# Neuronal thresholds and correlations in the peripheral vestibular system during rotation discrimination

**DOI:** 10.1101/343764

**Authors:** Courtney D. Garcia, Sheng Liu, Jean Laurens, Gregory C. DeAngelis, J. David Dickman, Dora E. Angelaki

## Abstract

Neuronal and behavioral thresholds were measured simultaneously as trained male macaques performed a yaw rotation discrimination task in darkness. When corrected to account for variations in neuronal direction preferences, neurons in the vestibular nuclei and semicircular canal afferents had discrimination thresholds that were only two-fold smaller than behavioral thresholds. There was no significant trial-by-trial correlation between neuronal activity and perceptual decisions, despite the presence of significant pair-wise noise correlations. The lack of choice-related activity during rotation discrimination contrasts with the robust correlations observed previously between brainstem neurons and choices during translation perception. These results suggest task-dependent differences in subcortical processing of vestibular signals, as well as how signals related to perceptual decisions may propagate back to early stages of sensory processing.

**SIGNIFICANCE STATEMENT:** This is the first ever simultaneous recordings of neural and behavioral thresholds during rotation discrimination. Its importance lies on the fact that the vestibular system provides an excellent model to probe origins of perception because directional selectivity signals are similar at many levels of processing, from afferents to cortex. The findings of similar neuronal and behavioral discrimination thresholds, significant inter-neuronal correlations, but lack of correlations between behavior and neuronal activity of both afferents and central brainstem neurons are intriguing and suggest task-dependent organization of early sensory areas.

## INTRODUCTION

Understanding how populations of sensory neurons represent information and how these populations are decoded to drive perception continues to be a topic of major interest in systems neuroscience. Specifying how much information is carried by individual neurons and how neural activity is correlated with perceptual decisions has been an important step in this endeavor (Parker and Newsome 1998; Nienborg et al. 2012). The vestibular system provides important opportunities for addressing these questions because neurons with similar stimulus selectivity can be studied from afferents to high-level association cortex (Laurens et al. 2017); this is not possible for other model sensory systems, such as visual motion and depth perception in primates.

In the vestibular system, previous studies have made direct comparisons between perception of translational motion (i.e., heading) and neural activity in cortex (Gu et al. 2007; Chen et al. 2013) and subcortical areas (vestibular nuclei: Liu et al. 2013a,b; otolith afferents: Yu et al. 2015) to explore how neural signals correlate with both the sensory stimulus and the binary perceptual decision. We found that neuronal discrimination thresholds of individual otolith afferents and neurons in the vestibular nuclei (VN) resemble those in cortical areas, and that the most sensitive cells have thresholds that are only two-fold greater than perceptual thresholds (Yu et al. 2015).

Previous studies have also characterized trial-by-trial correlations between neural activity and animals’ choices in a perceptual discrimination task. Choice-related signals are often quantified as ‘choice probabilities’ (CPs), as described by Britten et al (1996). CPs above chance have now been reported in many cortical areas and multiple sensory systems (Cumming and Nienborg 2016, Nienborg et al. 2012). In the vestibular system, Liu et al. (2013a) reported robust correlations between VN activity and perceptual decisions in a fine heading discrimination task. Remarkably, large CPs were found in some VN neurons that are selective to translation, but not in otolith afferents or other VN neurons that only represent the net gravito-inertial acceleration (Yu et al. 2015). We reasoned that response fluctuations of the latter groups of neurons are not linked to behavior because these neurons fail to discriminate self-motion from changes in orientation relative to gravity (Liu et al. 2013a).

By comparison, little is currently known about vestibular rotation perception and how it is shaped by early sensory processing. In contrast to translation, which has been studied in multiple brain areas (Liu et al. 2013a,b; Chen et al. 2013; Gu et al. 2007; Yu et al. 2015), there are no reports comparing simultaneously-recorded neural and behavioral measures of sensitivity to rotation. Instead, neuronal thresholds for yaw rotation have only been measured for horizontal canal afferents (Sadeghi et al. 2007a,b; Yu et al. 2014) and VN neurons (Massot et al. 2011; Yu et al. 2014) in response to sinusoidal rotation stimuli presented to untrained animals. Macaque neuronal thresholds for rotation were instead compared with human perceptual thresholds (Chaudhuri et al. 2013; Grabherr et al. 2008; Haburcakova et al. 2012; Kolev and Nicoucar 2014; Lim and Merfeld 2012; Mallery et al. 2010; Seemungal et al. 2004; 2009; Soyka et al. 2012) to assess whether neural activity was sufficient to account for perceptual sensitivity. However, macaques and humans may not have identical behavioral sensitivity, perceptual thresholds may vary substantially across subjects (e.g., Valko et al. 2012; Liu et al. 2013a, see Sup Fig 2C), and variations in vibration among motion platforms may also affect threshold measurements (Merfeld 2011; Lim and Merfeld 2012). Thus, it is critical to compare simultaneous measures of neuronal and behavioral performance in the same subjects.

Here, for the first time, we compare simultaneously-measured neuronal and perceptual rotation discrimination thresholds, and we search for correlations between neural activity and perceptual decisions during rotation discrimination. We focus on peripheral vestibular areas, including afferents from the semi-circular canals and neurons in the vestibular nuclei, as the response properties in these areas place important constraints on subsequent stages of neural processing and behavior. By comparing our findings to previous data obtained during fine discrimination heading, these experiments allow us to examine how the links between activity of single neurons and perception depend on the nature of the task and the origins of the signals.

## METHODS

### Subjects and equipment

Four male rhesus monkeys (*Macaca mulatta*) were implanted chronically with a circular head-restraint ring and recording grid for controlled stereotaxic electrode penetrations. When head-fixed using this restraint ring, the horizontal stereotaxic plane was parallel to the earth-horizontal plane (elevating the horizontal semicircular canals by approximately 14°; Blanks et al. 1985; Reisine et al. 1988; Haque et al. 2004). Eye movements were monitored either using chronically implanted eye coils (Angelaki and Dickman 2000) or a video eye tracker (ISCAN). All surgical and experimental procedures were approved by the Institutional Animal Care and Use Committee at Baylor College of Medicine and were performed in accordance with institutional and NIH guidelines.

Experiments were performed in head-fixed animals positioned on a 6 degree-of-freedom hexapod platform (MOOG6DOF2000E) that delivered the motion stimuli. Visual targets were rear-projected onto a tangent screen at a viewing distance of 32.5 cm by a video projector (Christie Digital, model MIRAGE S+3K). Electrodes were advanced inside 26-gauge transdural guide tubes using a remote controlled microdrive (FHC). In initial recordings, we identified the abducens and vestibular nuclei using single electrodes (Frederick Haer, tip diameter 3 μm, impedance 5-7 MΩ) following procedures similar to those used in previous studies (Angelaki and Dickman 2000; Meng et al. 2005; Meng and Angelaki 2006).

Once the area around the VN was mapped, we used linear array electrodes (Plexon U-Probe; 16 contacts, 100 μm apart) in two animals, monkey I and monkey T, to collect data in 68 recording sessions (Fig. 1). Note that linear array electrodes allowed us to sample VN neurons in a relatively unbiased manner, without prescreening (see Discussion).

**Fig. 1.**
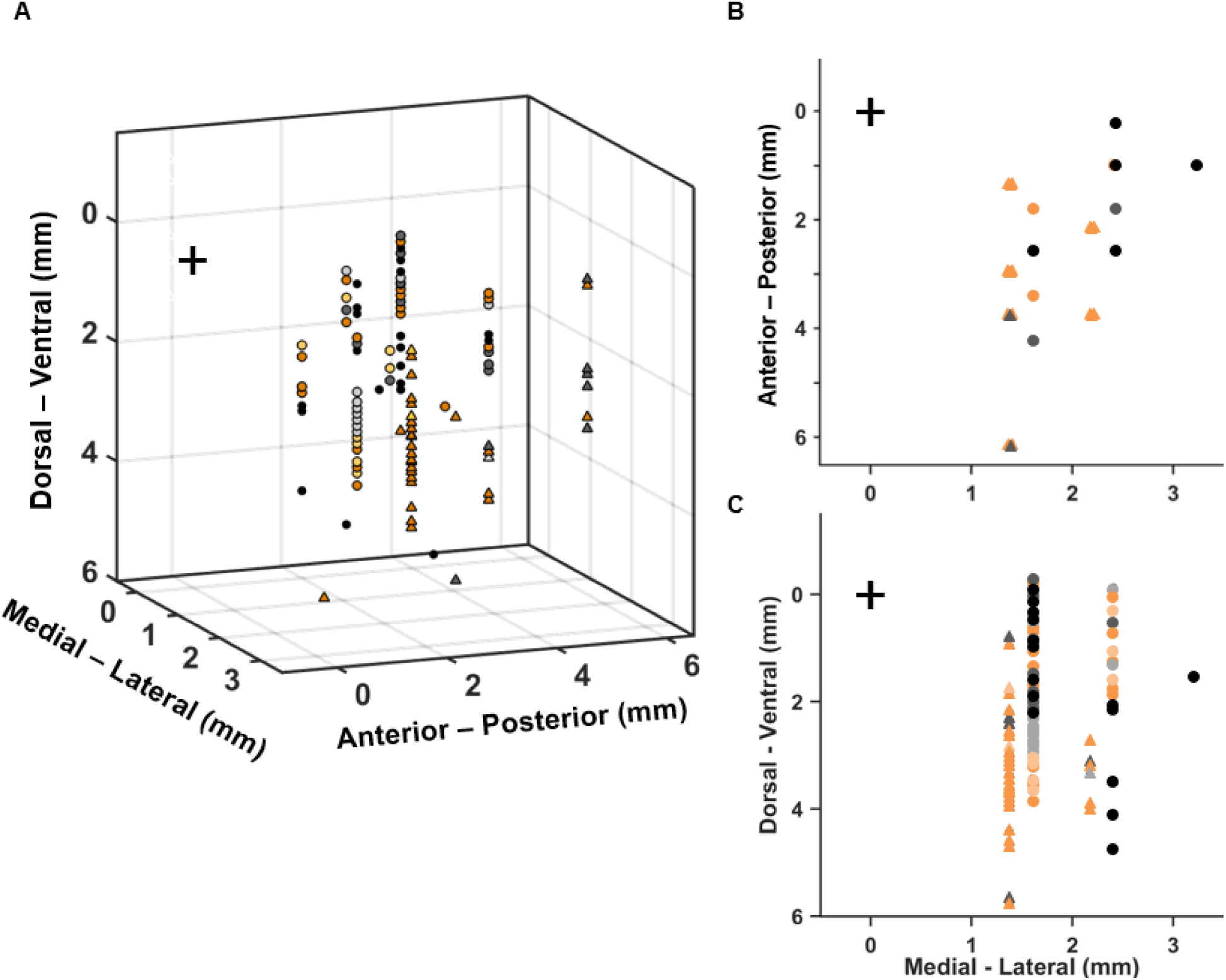
Reconstruction of VN recording sites. **(*A*)** Three dimensional illustration of recording site locations. **(*B*)** Recording locations projected onto the horizontal stereotaxic plane. **(*C*)** Recording locations projected onto the coronal plane. Each symbol represents a recorded cell, classified as *Uni-directional hybrid* (filled black, n = 29), *Uni-directional facilitated* (dark gray, n = 21), *Uni-directional suppressed* (light gray, n = 12), *Bi-directional facilitated* (dark orange, n = 58), or *Bi-directional suppressed* (light orange, n = 11). Cell locations as plotted as if all cells were recorded from the right brainstem (although data from both hemispheres are included). Symbol shapes denote different animals: monkey I (n = 82, ○) and monkey T (n = 49, Δ). The location of the abducens nucleus was taken as the origin of the coordinate system (cross at [0,0]).

In addition, recordings were obtained from horizontal canal afferents in three animals, monkey I, monkey H, and monkey Y, using single tungsten epoxy-coated electrodes (impedance 18-20 MΩ) following established procedures of localization and identification (Yu et al. 2012; 2015). Semicircular canal afferents were encountered directly below the cerebellum, as they entered the brain through the internal auditory meatus (see Haque et al. 2004; Yu et al 2012). Upon isolation, each fiber was first tested with 0.5 Hz, ±7° yaw, pitch and roll sinusoidal rotations to identify horizontal canal afferents (Haque et al. 2004) for further testing in the yaw discrimination task.

Prior to recording experiments, the two animals used for VN recordings were trained to pursue visual targets and to perform a two-alternative forced choice rotational motion discrimination task around psychophysical threshold (Fig. 3A). In this task, animals were trained (until they reached asymptotic performance) to report their perceived rotation direction (rightward vs. leftward). Each trial started when the animal fixated a visual target located straight in front of the animal’s head for 400 ms. Upon satisfactory fixation within a 3° fixation window, the target was turned off and the motion stimulus started. On each trial, the velocity profile followed a 0.7s trapezoidal waveform (100 ms acceleration, 500 ms constant velocity, 100 ms deceleration). White noise was played from speakers within the room to mask any auditory cues from the motion platform. At motion’s end there was a 400 ms delay, then the animal was required to make a saccade to one of two visual targets, located 10 deg to the right and left of straight ahead, to report his perceived motion direction. When the answer was correct, the animal received a juice reward. For the ambiguous yaw rotation velocity (0°/s), random delivery of reward was given on half of the trials.

### Experimental protocol

Once the U-Probe was lowered into the brainstem, the tissue was allowed to settle for ~30-60 minutes, and then the following experimental protocols were delivered:

1. Yaw rotation during visual fixation (‘yaw fixation task’): Animals were rotated about the yaw axis (Gaussian velocity profile, 12° amplitude, 1-s duration, 10 repetitions of both leftward and rightward rotations), while maintaining visual fixation on a head-fixed target.
2. Horizontal and vertical pursuit eye movements: Animals pursued a target moving across the screen in four cardinal directions: down, up, left, and right (trapezoidal velocity profile, 7° amplitude, 2-s duration, five repetitions/direction).
3. Three-dimensional sinusoidal stimuli: Animals were rotated in darkness using 0.5 Hz yaw, pitch and roll rotations (±21.9°/s peak velocity, 10 cycles), as well as translation along the three cardinal axes (interaural and naso-occipital axes: ±31.4 cm/s peak velocity; vertical axis: ±21.9 cm/s, 20 cycles).
4. Rotation discrimination task: In a block of randomly interleaved trials, the following set of peak angular velocities (trapezoidal velocity profile) was presented: 0°/s, ±0.83°/s, ±1.67°/s, ±3.33°/s, ±6.67°/s, and ±13.33°/s. The range of speeds was selected in order to: i) include stimuli small enough (e.g., 0.5 – 1 °/s) to accurately measure behavioral thresholds, and ii) include stimuli large enough (e.g. ±13.33 °/s) to measure neuronal tuning and thresholds for neurons that are not highly sensitive. Animals wore shutter glasses which completely obstructed their vision during platform motion. Animals were required to return their gaze to within the fixation window by the end of the motion stimulus, and generally tended to maintain their eye position around straight ahead during platform motion. Trials were aborted if the eye was not within ±1.5° of the fixation target when the shutter glasses opened at the end of the trial.

When recording from vestibular afferents, only protocols 3 and 4 were delivered.

### Data analyses

#### Analysis of VN neuronal activity

Neural signals were processed and spike-sorted off-line using the Plexon Offline Sorter (version x64 V3). We analyzed data from any well-isolated neuron, regardless of responsiveness to the stimuli we used, a notable difference compared to previous VN studies, which all pre-selected neurons according to their response characteristics. Our approach also allowed us to record from many VN cells that were silent in the absence of motion, another difference from previous studies.

To test whether neurons were responsive to smooth pursuit eye movements, we measured each neuron’s average firing rate during each pursuit trial (n = 5 repetitions for each of the 4 pursuit directions). We used a one-way ANOVA (3, 16 dof) to compare the responses across the four directions. Eleven of the 69 neurons recorded were significantly tuned to the direction of smooth pursuit at *P* < 0.01, and were excluded from our sample. Thus, we focused on VN cells without significant eye movement sensitivity.

To test whether neurons were responsive to sinusoidal motion (translation or rotation about the three cardinal axes), we computed the sinusoidal response gain and phase for each motion type by fitting a rectified sinusoid to the temporal response profile (firing rate vs. time) (Yakusheva et al. 2007). We assessed the significance of response modulation for each stimulus direction using a shuffling procedure (n = 1000 samples), in which the entire motion stimulus sequence was divided into individual cycles (n = 20 cycles, 2s duration each). The neuron’s response was shifted circularly within each cycle, and the gain and phase of the average response across all cycles was recomputed. A neuron was considered to be significantly tuned (*P* < 0.01) to one motion type if the average gain in the un-shuffled data was greater than the 99th percentile of the shuffled data. We classified neurons as ‘rotation only’ if they were significantly tuned to at least one rotation direction but no translation direction, and ‘convergent’ if they were significantly tuned to at least one rotation direction and one translation direction.

We also computed each VN neuron’s three-dimensional preferred direction (PD) of rotation and response gain (spk/s/°/s) based on responses to 0.5 Hz sinusoidal rotational motion about the yaw (z), pitch (y) and roll (x) axes, assuming cosine tuning. PDs are three-dimensional vectors [PD_x_, PD_y_, PD_z_] representing the inferred axis around which rotational responses should be maximal (Laurens et al 2015). In order to characterize cells that responded preferentially to yaw rotational motion, we computed the absolute angle between the PD and the yaw axis (referred to as the ‘difference angle’): α = acos([PDx, PDy, PDz] [0 0 1]), where ‘·’ represents the vectorial dot product.

We also quantified responsiveness to yaw rotation during visual fixation as follows: the neuron’s baseline firing rate was computed during a window of 400 ms preceding each yaw rotation stimulus, and this baseline response was compared to the neuron’s firing rate in a window of 400 ms centered on each trial’s peak stimulus velocity (Wilcoxon rank sum test). This analysis was performed independently for rotations to the right and to the left. Neurons were considered significantly responsive to leftward or rightward rotation if the Wilcoxon test yielded a p-value < 0.05 for the respective yaw directions.

For this group of responsive VN cells, spontaneous firing rate was calculated from the 400 ms preceding each yaw stimulus. Modulation amplitude (spikes/s) was calculated as the absolute difference between peak firing rate within the trial and spontaneous activity. Furthermore, cells with significant responsiveness were classified as ‘facilitated’ or ‘suppressed’ for each yaw direction.

These comparisons allowed us to distinguish VN neurons as either ‘unidirectional’ or ‘bi-directional’ (Fig. 2 and Table 1). Uni-directional neurons were of three types: The first two types included ‘facilitated’ or ‘suppressed’ during only one direction of rotation (did not pass the ‘responsive’ criterion in the other yaw direction). The third type, which we called ‘hybrid’, was responsive to both yaw directions, facilitated and suppressed in opposite directions. Bi-directional neurons were also responsive to both yaw directions, but were of the same modulation type, i.e., either both facilitated or both suppressed.

**Fig. 2.**
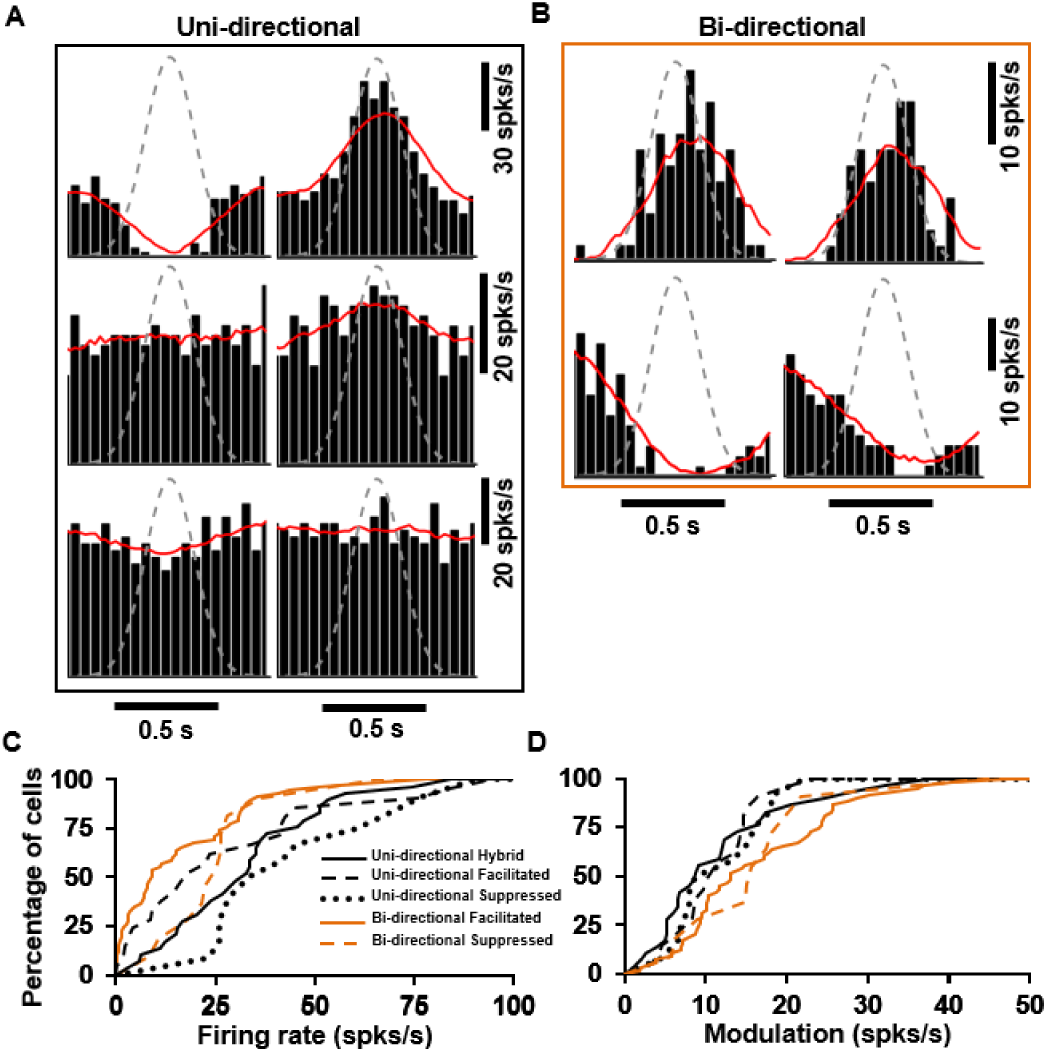
Temporal profiles of responses to yaw rotation in the VN. **(A), (B)** Example PSTHs illustrating temporal profiles observed in response yaw rotation during visual fixation of a central target. Left panel of each pair: leftward rotation; Right panels: rightward rotation. Gray dashed lines: stimulus speed profile. Red lines: best-fit Gaussian function. *(A, top row):* Example neuron showing *Uni-directional hybrid* response, in which firing rate increases for one direction of rotation and decreases for the other. *(A, bottom two rows):* Examples of *Uni-directional* responses, in which firing rate either increases (*middle row*) or decreases (*bottom row*) during rotation in one direction, with no response to rotation in the opposite direction. *(B):* Two example neurons showing *Bi-directional* response modulation, in which firing rate either increases *(top)* or decreases *(bottom)* during rotation in both directions. ***(C, D)*** Cumulative distributions of resting firing rate (C) and response modulation amplitude (D). Data are shown separately for different cell types: *Uni-directional hybrid* (black solid lines, n = 29 cells), *Uni-directional facilitated* (black dashed lines, n = 21 cells), *Unidirectional suppressed* (black dotted line, n = 12 cells), *Bi-directional facilitated* (orange solid lines, n = 58 cells), and *Bi-directional suppressed* (orange dashed lines, n = 11 cells).

**Table 1.**
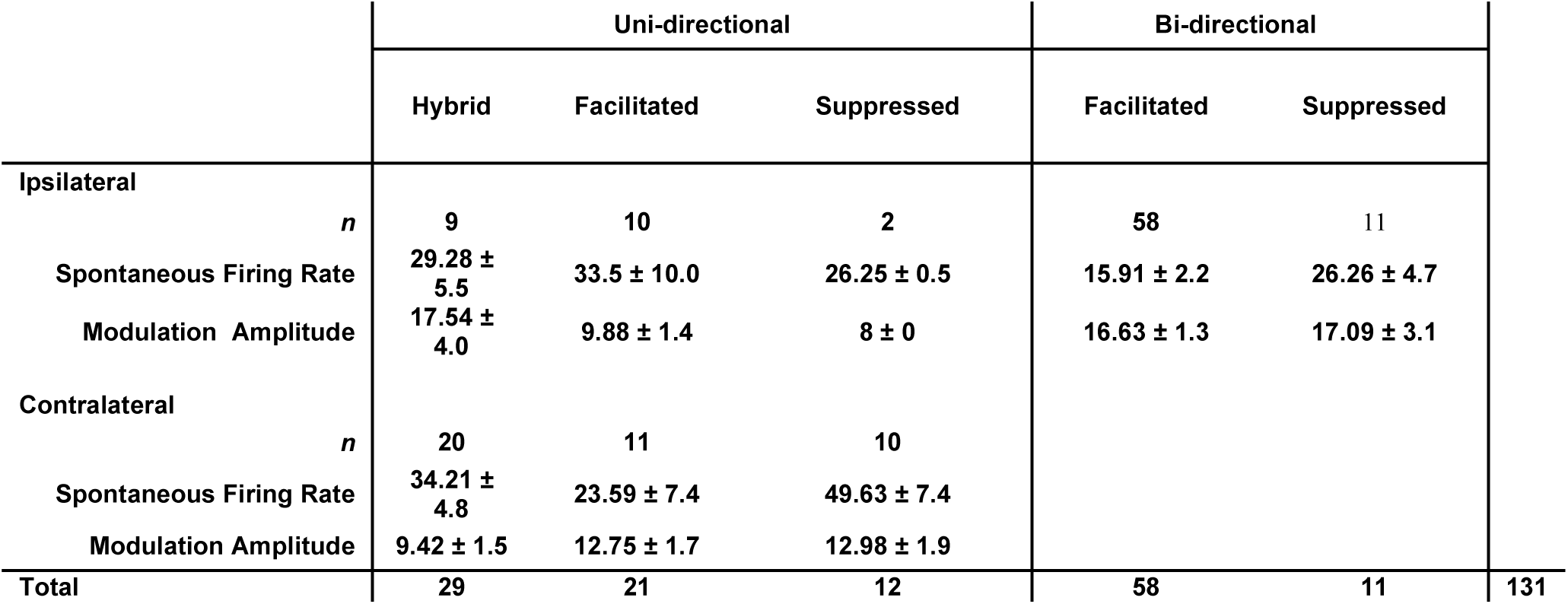
Classification of VN neurons based on response modulation to ipsilateral or contralateral yaw rotation (relative to the recording side; monkey I, n = 82; monkey T, n = 49). For neurons with Uni-directional hybrid modulation, ipsilateral/contralateral refers to firing rate increases. Spontaneous firing rate (mean ± s.e.m.) was calculated from the 400 ms preceding each stimulus presentation (yaw fixation task, Gaussian velocity profile). Modulation amplitude was computed as the absolute difference between spontaneous firing rate and maximum or minimum firing rate, for facilitated or suppressed responses respectively, within the trial.

For all VN neurons responsive to rotation in the yaw fixation task, only those whose isolation was maintained during at least 10 repetitions of the rotation discrimination task were further analyzed as follows. First, we computed neuronal discrimination thresholds during the rotation discrimination task. Spike counts during the middle 500 ms of the stimulus, when velocity was constant, were used to perform a receiver operating characteristic (ROC) analysis (see Gu et al. 2007; Liu et al. 2013a for details). This analysis computed the ability of an ideal observer to discriminate between two oppositely-directed yaw rotations with the same peak speed (e.g., −13.33°/s versus +13.33°/s; Fig. 3C, E, ⍰) based exclusively on the firing rate of the recorded neuron and a supposed ‘anti-neuron’ with opposite tuning (Britten et al. 1992). ROC values were plotted as a function of yaw velocity, resulting in a neurometric function that was fit with a cumulative Gaussian curve (Fig. 3C, E, solid line). Neuronal threshold was defined as the standard deviation of the Gaussian fit. For a minority of insensitive neurons (n = 5), threshold values were restricted to an upper limit of 300°/s.

**Fig 3.**
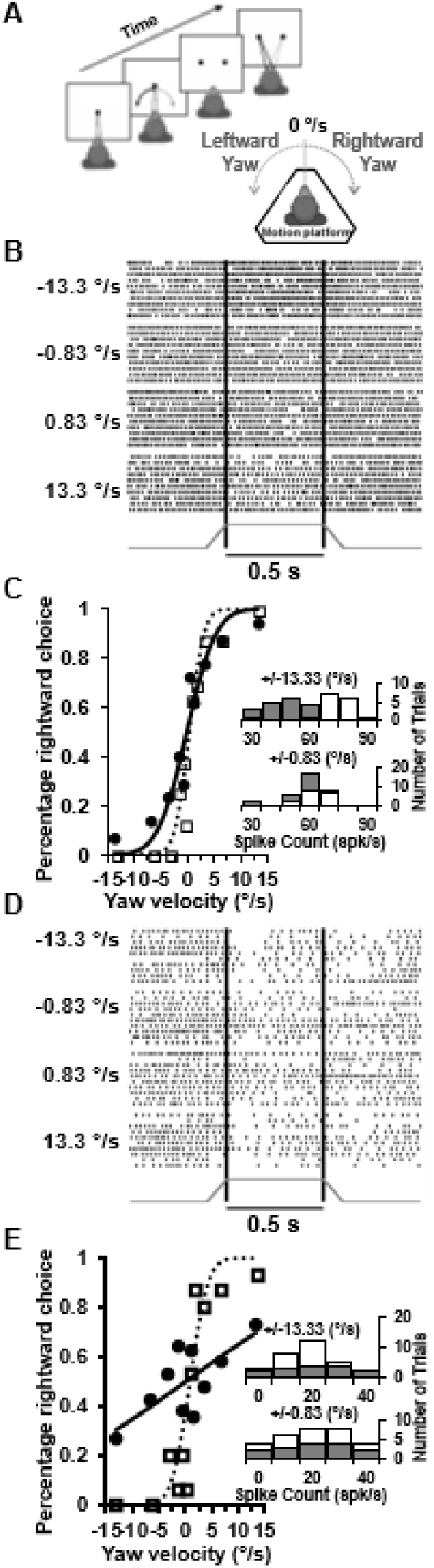
Rotation discrimination task and analysis of responses of two example VN neurons. ***(A)*** Schematic illustration of yaw rotation discrimination task. ***(B, D)*** Raster plots of neural responses to leftward and rightward yaw rotation at two speeds. For the uni-directional VN neuron *(B),* firing rate in a 500 ms window (constant velocity period) increased for contralateral rotation (leftward, negative numbers) and decreased for ipsilateral (rightward, positive numbers) rotation. For the example bi-directional neuron (*D),* firing rate decreased for both rotation directions. ***(C, E)*** Neurometric functions (and solid curves) for the example VN neurons. The corresponding psychometric functions are also shown (and dotted curves). Insets distributions of spike counts for ipsilateral (filled bars) and contralateral (open bars) rotations (for the same rotation velocities as in B, D).

Second, behavioral performance was quantified by plotting the proportion of rightward choices as a function of peak yaw velocity (Fig. 3C, E, ⍰) and fitting the data with a cumulative Gaussian function (Fig. 3C, E, dotted line). The SD of the cumulative Gaussian fit was defined as the behavioral threshold (Wichmann et al. 2001). For each session, neuronal and behavioral thresholds were calculated from data gathered simultaneously, allowing for a direct comparison between them. We divided the psychophysical thresholds by √2 to account for the behavioral task being conducted as a one interval task (Hillis et al. 2004), while the neurometric functions were computed by comparing two different distributions (the neuron and its hypothetical anti-neuron). Geometric means and SDs were calculated by computing the mean and SD of the logarithm of the psychophysical and neuronal thresholds.

Third, we used ROC analysis to compute choice probability (CP), which quantifies the relationship between neural responses and choices, independent of the stimulus. The data were normalized using balanced Z-scoring (Kang et al. 2012) for each stimulus with at least 3 choices in each direction and were then combined into a single pair of distributions corresponding to preferred and null choices, where a preferred choice is a choice in favor of the neuron’s preferred yaw rotation direction. The CP was determined by performing ROC analysis on this pair of distributions, and statistical significance was based on bootstrapping to determine whether a particular CP was significantly different, *P* < 0.05, from the chance level of 0.5.

Fourth, we used the number of spikes recorded during the middle 500 ms of the stimulus (constant velocity phase) to measure ‘signal’ (r_signal_) and ‘noise’ (*r*_noise_) correlations for pairs of VN neurons recorded simultaneously during the yaw rotation discrimination task. The signal correlation, r_signal_, was computed as the Pearson correlation coefficient (which ranges from −1 to 1) between tuning curves (mean firing rate as a function of stimulus velocity) measured during the discrimination task for each pair of VN neurons (Liu et al. 2013a). Noise correlation, r_noise,_ was computed as the Pearson correlation coefficient of the z-scored individual trial responses of each pair of neurons (Zohary et al. 1994). Responses were re-normalized (z-scored) in blocks of 20 trials to remove slow fluctuations in firing rate that could arise from changes in cognitive or physiological state (Zohary et al. 1994; Lu et al. 2013a).

Based on neuronal thresholds measured during yaw rotation, we extrapolated the threshold along each neuron’s preferred direction (Yu et al. 2015; Jamali et al. 2013) by using the formula: (threshold along the preferred direction) = (threshold during yaw rotation)*cos(α), where α is the angle between the head’s yaw axis and the neurons’ preferred direction. We then computed the geometric mean across the population. Neurons with a difference angle α ≥ 89°(n = 2/58) were excluded from this computation because cos(90) = 0.

#### Analysis of horizontal semicircular canal afferent activity

Computation of neuronal thresholds and CPs from responses of horizontal canal afferents during the rotation discrimination task was identical to that described above for the vestibular nuclei. Discharge regularity was quantified from the spontaneous activity of canal afferents. The distribution of ISIs recorded during spontaneous activity was used to compute the coefficient of variation (CV), CV = σ_ISI_/μ_ISI_, where σ_ISI_ and μ_ISI_ denote the SD and mean of the ISI distribution. A normalized measure, CV*, was used to classify canal afferents as regular (CV* < 0.15) or irregular (CV* ≥ 0.15), because CV varies with the mean ISI (Haque et al. 2004; Sadeghi et al. 2007a,b; Goldberg et al. 1984).

## RESULTS

### Types of responses found in the VN

We recorded from the vestibular nuclei of two macaques, sampling an area posterior and lateral to the abducens nucleus (Fig. 1). Out of 183 neurons with well-isolated spikes (from 68 U-Probe recording sessions), 131 cells showed significant responsiveness to yaw rotation (see Methods). Of these modulated cells, a total of 62 VN neurons were significantly tuned for the *direction* of yaw rotation (hereafter, ***‘uni*-directional*’ VN*** cells) and included three cell types: one type, called ‘hybrid’, increased firing rate during rotation in one direction, while decreasing firing rate during rotation in the opposite direction (n = 29; Fig. 2A, top row; Table 1). Other subgroups of unidirectional neurons showed increased or decreased their firing rates during rotation in only one direction, with no significant response change for the opposite direction (‘unidirectional facilitated’ and ‘uni-directional suppressed’ cells, respectively, n = 33; Fig. 2A, bottom two rows; Table 1). Perhaps surprisingly, we also identified a large group (n = 69) of VN neurons that increased or decreased their firing rates during rotation in *both* directions ***(‘bi-directional’ VN*** cells; Fig. 2B; Table 1). Such bi-directional response types have previously been found in reduced animal preparations (Type III classification of Duensing and Schaeffer 1958), but have never before been described in the VN of alert animals (see Discussion).

Unlike in previous studies, the present recordings were performed using linear electrode arrays and we recorded from neurons without any pre-screening of responses. The mean resting firing rate for bi-directional VN cells is significantly lower than that for uni-directional VN cells (*P* = 1.38 × 10^−5^, Wilcoxon rank sum test; Fig. 2C, Table 1). Some cells have resting firing rates of 0 spikes/s; these cells may not have been included in previous single-electrode recording studies. Bi-directional VN cells also exhibited significantly greater response modulation amplitudes than uni-directional VN cells (*P* = 2.4 × 10^−3^, Wilcoxon rank sum test; Fig. 2D, Table 1). We found no significant differences in modulation amplitude among the different subtypes within the unidirectional and bi-directional cell groups. Despite these differences, uni-directional and bi-directional cells appeared intermingled throughout the recorded area (Fig. 1).

### Rotation discrimination: VN neuronal thresholds

Of the 131 VN neurons with significant response to yaw rotation during visual fixation (‘yaw fixation’ task), we searched off-line for those with well-isolated spikes throughout at least 10 repetitions of the yaw discrimination task (Fig. 3A). A subgroup of 58 cells met these criteria; they modulated during yaw fixation but did not respond during either pursuit eye movements (see Methods) nor during fast phases of nystagmus elicited by sinusoidal rotation stimuli. Example neuronal and behavioral responses are illustrated in Fig. 3B,C for a uni-directional VN cell with hybrid response modulation. Mean spike counts during the middle 500 ms of constant velocity rotation were used to compute a neurometric function (Fig. 3C, filled symbols) that was fit with a cumulative Gaussian (Fig. 3C, solid curve) to extract a neuronal threshold, which was 4.3°/s for this neuron. These data were then compared with the animal’s psychometric function (Fig. 3C, open symbols and dotted curve), which yielded a behavioral threshold of 1.5°/s. Thus, the example uni-directional hybrid neuron in Fig. 3C was only modestly less sensitive than the animal. An analogous set of data are shown in Fig. 3D,E for a bidirectional VN cell. For this cell, the neuronal threshold of 26.1°/s was much greater than the behavioral threshold of 1.9°/s.

A summary of all simultaneously-recorded psychometric and neurometric functions is shown in Fig. 4A,B, along with the corresponding cumulative Gaussian fits. VN neuronal thresholds had a geometric mean of 31.9°/s (95% confidence interval: [22.7 - 44.9]), as compared to the geometric mean behavioral threshold of 2.3°/s [95% CI: 2 – 2.5]. When separated into uni-directional and bi-directional cell types, geometric mean neuronal thresholds were 11.9°/s [95% CI: 7.3 – 19.3] and 56.1°/s [95% CI: 34.9 – 90], respectively, and this difference was significant (*P* = 1.56 × 10^−5^, Wilcoxon rank sum test). Across all neurons, VN neuronal thresholds and goodness-of-fit of the cumulative Gaussian were strongly correlated (Spearman rank correlation: ρ = 0.89, *P* = 1.07 × 10^−20^), such that greater thresholds were typically associated with lower R^2^ values (Fig. 4B, inset). This was expected because high neuronal thresholds correspond to flat neurometric functions. The greater neuronal thresholds of bidirectional neurons suggest that these cells do not provide much useful information for discriminating the direction of yaw rotation.

**Fig 4.**
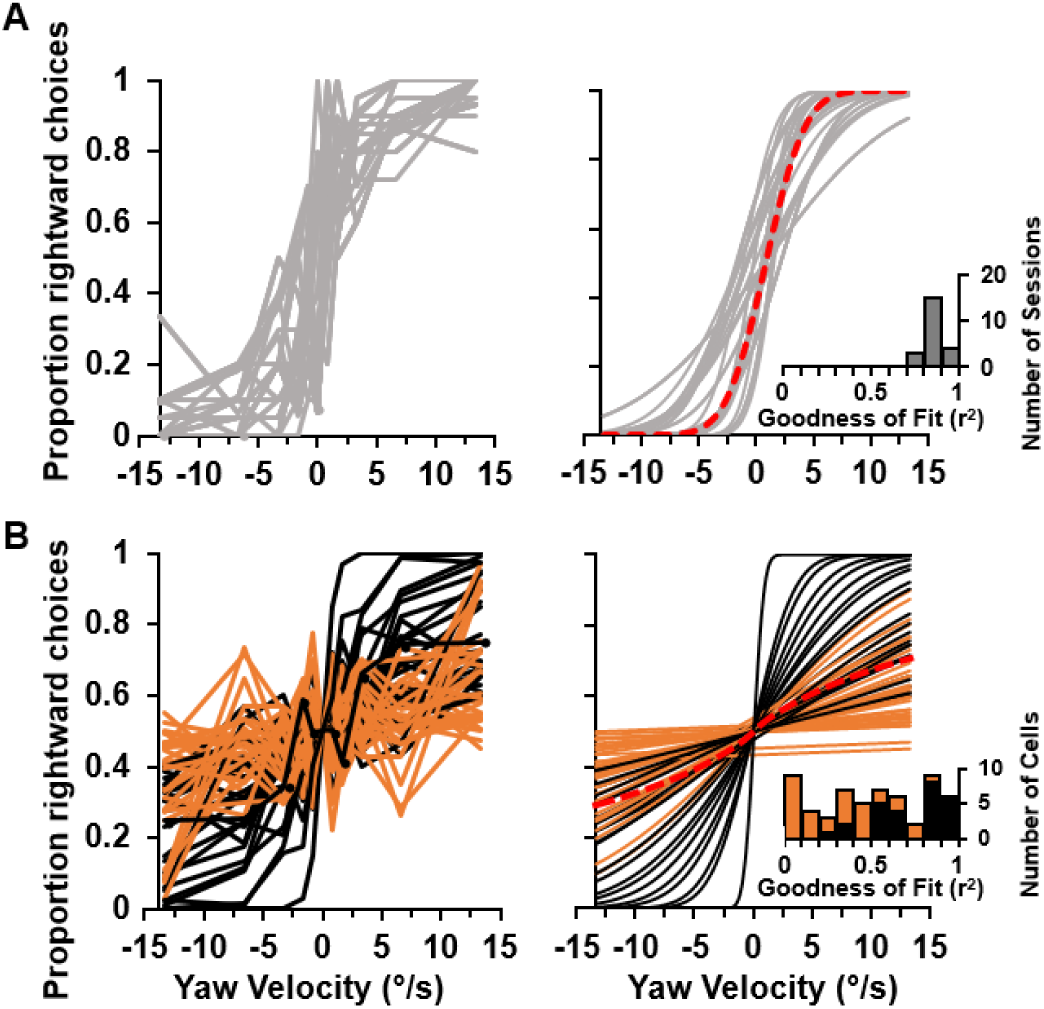
Summary of psychometric and neurometric functions. ***(A)*** Psychometric data (left) and corresponding cumulative Gaussian fits (right) from 22 recording sessions. Inset: R^2^ distribution of Gaussian fits (right). ***(B)*** Neurometric data (left) and corresponding fits for uni-directional (black, n = 26) and bi-directional (orange, n = 32) VN neurons. Red dashed lines: mean psychometric and neurometric fits. Inset: R^2^ distribution of Gaussian fits (right).

The relationship between neuronal and psychophysical thresholds is summarized in Fig. 5A. For all but one VN cell, the neuronal threshold was greater than the psychophysical threshold (Fig. 5A, blue and green symbols). VN neurons were further categorized as ‘convergent’ or ‘rotation-only’ based on their responses to sinusoidal translation stimuli (see Methods), and these types differed in their sensitivity. The ratio of neuronal:psychophysical thresholds had a geometric mean of 17 [95% CI: 12.4 – 23.3] for convergent cells and 1.8 [95% CI: 0.2 −16.6] for rotation-only VN cells (Fig. 5A, blue and green symbols, respectively), and this difference was significant (*P* = 5.6 × 10^−3^, Wilcoxon rank sum test). The geometric mean neuronal threshold for convergent VN cells was 38.5°/s [95% CI: 28.2 – 52.6, n = 53], whereas the geometric mean threshold for rotation-only neurons was nine-fold smaller at 4.3°/s [95% CI: 0.8 - 22, n = 5] (*P* = 1.6 × 10^−3^, Wilcoxon rank sum test).

**Fig. 5.**
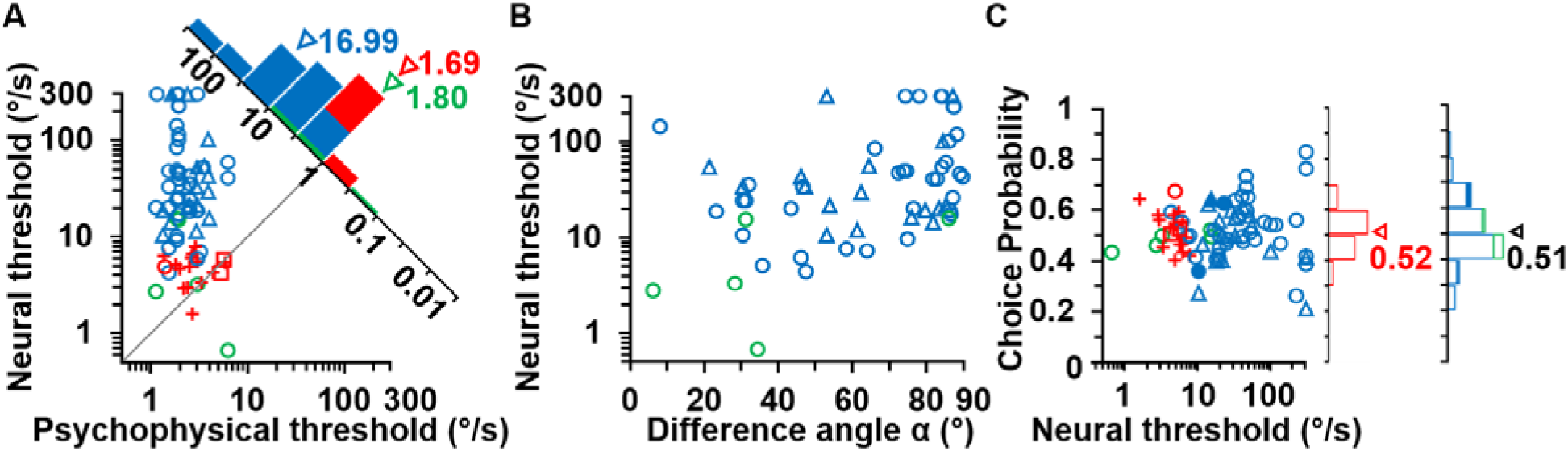
Summary of neuronal thresholds and choice probabilities. **(*A*)** Comparison of neuronal and psychophysical thresholds. The diagonal histogram shows the distribution of neuronal-to-psychophysical threshold ratios. Arrowheads illustrate geometric means. Psychophysical thresholds are divided by √2 for comparison to neuronal thresholds computed using the neuron/anti-neuron construct. **(*B*)** Neural thresholds are correlated with the difference angle, α, computed as the absolute difference between the yaw rotation axis and each neuron’s 3D preferred direction. ***(C)*** Choice Probabilities are plotted against neuronal thresholds. Filled and open symbols/bars indicate significant and nonsignificant CPs, respectively. Marginal histograms show distributions of CP and arrowheads show mean values. Data shown are for horizontal semicircular canal afferents (red, n = 18), convergent VN neurons (blue, n = 53), and rotation-only VN neurons (green, n = 5). Symbol shapes denote different animals: monkey I (VN n = 41, horizontal canal afferent n = 1, ○), monkey T (n = 17, Δ), monkey Y (n = 15, **+**), and monkey H (n = 2, ⍯).

A possible explanation for the greater thresholds of convergent VN neurons is the fact that their preferred directions are broadly distributed in 3D space (Dickman and Angelaki 2002). In contrast, the preferred directions of rotation-only cells tend to cluster around the semicircular canal axes, similar to canal afferents (Dickman and Angelaki 2002). To investigate whether the difference in sensitivity between convergent and rotation-only VN neurons may be due to their PDs, we computed the difference angle, α (°), as the absolute difference between the yaw rotation axis and each neuron’s 3D preferred direction of rotation (the latter was computed from responses to 0.5 Hz sinusoidal yaw, pitch and roll stimuli; see Methods). As expected, we found a significant correlation between neuronal thresholds and the difference angle (Spearman rank correlation: ρ = 0.41, *P* = 1.4 × 10^−3^, Fig. 5B), such that cells with preferred directions aligned with the yaw axis tended to be most sensitive. When the difference angle was split into 30° bins, neuronal thresholds had geometric means of 16.5°/s [95% CI: 1.9 - 140], 15.9°/s [95% CI: 8.5 – 29.3] and 48.1°/s [95% CI: 32.8 – 70.6] for sectors [0°, 30°], [31°, 60°], and [61°, 90°], respectively (ANOVA, *P* = 0.0056).

The few rotation-only cells in our sample do not appear to form a separate group in the scatter plot of Fig. 5B, which was supported by a non-significant interaction term in an analysis of covariance (ANCOVA *P = 0.75*). Among the rotation-only neurons, only one cell exhibited a difference angle of nearly 90°, whereas the rest had PDs close to the yaw axis (Fig. 5B, green). This rotation-only neuron with a large difference angle was classified as a bi-directional neuron (with increasing firing rate) and had a neuronal threshold of 15.6°/s, whereas all other rotation-only cells were classified as unidirectional and had small thresholds (geometric mean of 3.1°/s [95% CI: 0.4 – 23.3]). When thresholds were extrapolated for the preferred direction of each neuron (using an inverse cosine function; see Methods), rotation discrimination thresholds had a geometric mean of 8.5°/s [95% CI: 6 – 11.9] for VN neurons. When split into cell types, corrected geometric-mean VN thresholds were 5.9°/s [95% CI: 3.6 – 9.5] for all unidirectional VN cells, 2.3°/s [95% CI: 0.5 – 9.9] for rotation-only cells (n = 5), 7.0°/s [95% CI: 4.2 – 11.5] for convergent uni-directional cells (n = 18), and 11.5°/s [95% CI: 7.3 – 18.1] for convergent bi-directional neurons (n = 33).

### Choice Probabilities and correlated noise

Choice probability (CP) quantifies the relationship between neural responses and choices, independent of the stimulus. Previously, we reported large CPs in some VN neurons during a heading discrimination task, particularly for sensitive neurons that were also selectively tuned for translational motion (Liu et al. 2013a). In contrast, in the present experiments, we found that the average CP during the rotation discrimination task (0.51 ± 0.12, mean ± SD) was not significantly different from chance (*P* = 0.49, Wilcoxon signed rank test), and that CPs were not significantly correlated with neuronal thresholds across the population of VN neurons (Fig. 5C; Spearman rank correlation: ρ = 0.15, *P* = 0.27). Only 3 of 58 VN neurons had CPs that were significantly different from 0.5 (Fig. 5C, filled symbols), a proportion which is not greater than expected by chance.

It has also been suggested in previous studies that the magnitude of CPs is determined largely by the strength of correlated noise among neurons (Nienborg and Cumming2009;2010). We investigated whether the lack of significant CPs during rotation discrimination could be explained by weak noise correlations among VN neurons. We analyzed the trial-by-trial spike counts for all pairs of well-isolated VN neurons during the rotation discrimination task, as long as they were significantly responsive to yaw rotation, as shown for an example pair of neurons in Fig. 6A. The Pearson correlation coefficient of z-scored trial-by-trial responses measured noise correlation (*r*_noise_, Fig. 6C). In addition, we quantified the similarity of tuning by computing the Pearson correlation coefficient of the mean responses across rotation velocities (signal correlation, *r*_signal_; Fig. 6B).

**Fig. 6.**
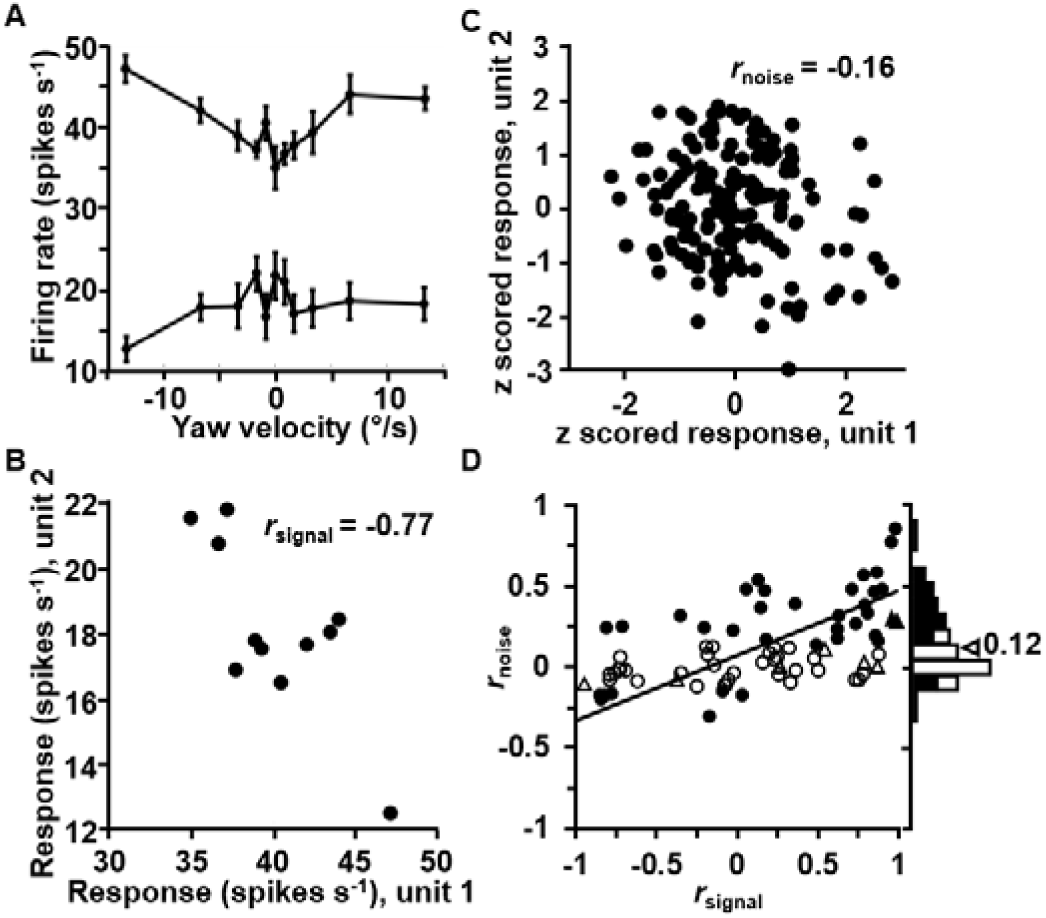
Summary of noise (*r*_noise_) and signal (*r*_signal_) correlations between pairs of VN neurons. **(*A*)** Tuning curves for an example pair of simultaneously recorded VN neurons. The tuning curves illustrate mean ± s.e.m. firing rates plotted as a function of stimulus peak velocity. **(*B*)** Comparison of the mean responses of the two example neurons across all rotational velocities; the corresponding Pearson correlation coefficient quantifies signal correlation, *r*_signal_ = −0.77 (*P* = 0.01). **(*C*)** Normalized responses from the same two neurons were weakly correlated across trials. Each datum represents *z* scored responses from one trial. The Pearson correlation coefficient of the data quantifies the noise correlation, *r*_noise_ = −0.16 (*P* = 0.03). **(*D*)** Scatter plot of *r*_noise_ versus *r*_signal_ for all pairs of simultaneously recorded VN neurons. Marginal distribution of *r*_noise_ is also shown (right). Arrowhead denotes the mean value of *r*_noise_. Filled symbols and bars denote *r*_noise_ values significantly different from zero (*P* < 0.05). Symbol shapes denote different animals: monkey I (n = 68, ○), monkey T (n = 8, Δ), and the solid line represents a type II linear regression fit.

Across our sample of pairwise recordings, noise correlations averaged 0.12 ± 0.03 (mean ± s.e.m., n = 76 pairs), and this value was significantly different from zero (*P* = 3.5 × 10^−5^; Wilcoxon signed rank test). Furthermore, we found a significant correlation between *r*_noise_ and *r*_signal_ (Spearman rank correlation, ρ = 0.56, *P* = 1.16 × 10^−7^; slope =0.40, intercept = 0.07; Fig. 6D), indicating that neurons with similar tuning tend to have larger positive noise correlation. For comparison, the average noise correlation measured during translation in our previous study was 0.004 ± 0.02 (mean ± s.e.m., n = 47 pairs; Liu et al. 2013a) and was not significantly different from zero (*P* = 0.7; Wilcoxon signed rank test). In that previous study, there was also a significant positive correlation between noise and signal correlations (Spearman rank correlation with *r*_signal_, ρ = 0.51, *P* = 2.84 × 10^−4^; slope = 0.528, intercept = −0.04). Thus, comparing the two studies, the mean *r*_noise_ was close to zero for translation and clearly above zero for rotation, but the relationship between noise and signal correlations was similar. The main difference between the two datasets seems to be that noise correlations tend to be negative for cell pairs with large negative signal correlations in the translation task, whereas noise correlations tend to be close to zero for large negative signal correlations during the rotation task. Importantly, the lack of significant CPs in the rotation discrimination task is not associated with a lack of correlated noise.

### Horizontal semicircular canal afferents

Data from horizontal semi-circular canal afferents in three animals (monkey I, n = 1; monkey H, n = 2, monkey Y, n = 15) were collected during an identical motion stimulus and rotation discrimination task. Canal afferents were classified as regular (CV* < 0.15; n = 10) or irregular (CV* > 0.15; n = 8) based upon the regularity of their discharge (see methods; Haque et al. 2004, Goldberg et al. 1984). Neuronal thresholds for regular and irregular canal afferents had geometric means of 4.6°/s [95% CI: 3.4 – 6.1] and 4.8°/s [95% CI: 3.4 – 6.5], respectively, and there was no significant dependence of neuronal thresholds on CV* (Spearman rank correlation: ρ = −0.02, *P* = 0.93), as Yu et al. 2015 found previously for otolith afferents. Additionally, we did not observe a significant difference between CPs for regular and irregular canal afferents (0.53 ± 0.09 and 0.51 ± 0.06, respectively, mean ± SD; *P* = 0.5, Wilcoxon signed rank test).

Comparing VN neurons and afferents, we find that the geometric mean threshold for canal afferents (4.7°/s [95% CI: 3.9 – 5.6], Fig. 5A,C) was not significantly different from that for VN rotation-only cells (*P* = 0.74, Wilcoxon rank sum test), but was significantly different from the other VN cell types (*P* = 1.59 × 10^−9^, Wilcoxon rank sum tests). These differences were also seen for neuronal-to-psychophysical threshold ratios, which had a geometric mean of 1.7 [95% CI: 1.3 – 2.2] for horizontal canal afferents (Fig. 5A histogram, red). As for VN neurons, the average CP of canal afferents (0.52 ± 0.08, mean ± SD) was not significantly different from chance (*P* = 0.32, Wilcoxon signed rank test).

We note that horizontal canal afferents all have the same preferred direction of rotation, which is determined by the orientation of the horizontal canals relative to the stereotaxic horizontal plane of the head. In macaques, this has been estimated to be ~15° (Blanks et al. 1985). Thus, we extrapolated the mean rotation threshold (measured during rotation in the stereotaxic horizontal plane) to rotation in the horizontal canal plane. This computed threshold for canal afferents (4.5°/s [95% CI: 3.7 – 5.5]) was significantly greater than the animals’ behavioral threshold (2.8°/s [95% CI: 2.2 – 3.4]; *P* = 0.0015, Wilcoxon rank sum test). It is remarkable that a single horizontal canal afferent is nearly as sensitive as the animal’s perception, given that there are many such canal afferents (see Discussion).

## DISCUSSION

The present findings demonstrate that: (1) The most sensitive semicircular canal afferents and VN neurons show rotation discrimination thresholds that are comparable to perceptual thresholds measured simultaneously in the same animal; (2) There are no significant correlations between perceptual decisions and the trial-by-trial activity of both VN neurons and canal afferents, despite the fact that there are significant interneuronal noise correlations among simultaneously recorded VN neurons. Each of these findings is discussed in more detail below.

### Comparison between neuronal and perceptual thresholds

We found that, although the average single neuron threshold is much greater than the behavioral threshold, the most sensitive VN neurons and all canal afferents have thresholds comparable to behavior. It is well established that the closer the stimulus to the preferred direction, the larger the response gain. Thus, as in previous studies (Gu et al., 2007; Yu et al., 2014; 2015), we found that neuronal thresholds depended on the absolute value of the difference between the yaw stimulus direction and the preferred 3D direction of the cell. Average neuronal thresholds corrected for the cells’ preferred direction were 5.9°/s for uni-directional VN neurons and 4.5°/s for horizontal canal afferents, as compared to the corresponding average behavioral thresholds of 2.2°/s and 2.8°/s, respectively. Notably, this is the first study to measure neuronal and behavioral thresholds simultaneously during rotation discrimination.

The finding that canal afferents and VN neurons can have thresholds comparable to behavior is consistent with results from a number of studies in both the vestibular and other sensory systems (Britten et al. 1992; Chen et al. 2013; Cohen and Newsome 2009; Gu et al. 2007, 2008). In contrast, Massot et al. (2011) and Sadeghi et al. (2007b) reported that VN and horizontal canal afferents had neural thresholds that were approximately an order of magnitude greater than behavioral thresholds. It is difficult to determine exactly which factors account for the large discrepancy between our findings and theirs because the stimuli used were not identical. However, it is worth noting that neuronal thresholds in our study are roughly similar to those reported by Massot et al. (2011) and Sadeghi et al. (2007b), suggesting that the comparisons to behavior may be a main source of the discrepancy. Cullen and colleagues (Massot et al. 2011, Sadeghi et al. 2007b) did not simultaneously measure behavior; rather, neuronal thresholds were compared with human perceptual thresholds (Grabherr et al. 2008; Haburcakova et al. 2012). Such cross-species comparisons are very difficult to interpret because one cannot assume that macaque and human thresholds are the same; furthermore, even within species, perceptual thresholds can vary substantially across subjects (e.g., Suppl. Fig. 2C of Liu et al. 2013a; Valko et al. 2012). In addition, the inherent stimulus-driven vibrations of motion platforms may also affect perceptual thresholds (Merfeld 2011; Lim and Merfeld 2012), complicating comparisons between different labs. Massot et al. (2011) also did not correct neuronal thresholds of VN neurons according to each cell’s preferred direction, which may also contribute to their conclusion that neurons are much less sensitive than behavior.

There are additional concerns with the conclusions of Massot et al. (2011), as well as Jamali et al. (2013), who quantified the pool size needed to convert macaque neuronal thresholds into human perceptual thresholds, assuming that population thresholds are proportional to the inverse square root of pool size. This assumption, however, ignores both correlated variability among neurons and the fact that population thresholds are determined by both encoding and decoding processes. Correlated noise cannot be assumed to be negligible in the VN, and thus should not be ignored because it may affect both the information carried by the population and decoding efficiency (Abbott and Dayan 1999; Averbeck et al. 2006; Ecker et al. 2011; Haefner et al. 2013; Sompolinsky et al. 2001; Shadlen et al. 1996). Furthermore, and perhaps most importantly, for the same neural pool size and correlated noise, different population thresholds may result depending on whether the decoder has full knowledge of these correlations or not (Haefner et al. 2013; Pitkow et al. 2015). Some noise correlations can be removed by an optimal decoder (Ecker et al. 2011), but the brain’s decoding of VN activity may not be optimal (Pitkow et al. 2015). Other noise correlations cannot be removed by any decoder, such that the total possible decoded information saturates regardless of the decoder (Moreno-Bote et al. 2014; Pitkow et al. 2015). Importantly, what is ultimately important for decoding and perception is not the pairwise correlations measured here (and in many other experimental studies), but rather the multidimensional covariance matrix, whose measurement is at present limited by technology. Estimating the covariance matrix of a population from small (e.g., pairwise) recordings is known to be very error-prone (Averbeck et al. 2006). Thus, although comparisons between single neuron and behavioral thresholds can be made when both are recorded simultaneously, extrapolation to population thresholds is faced with numerous problems and uncontrolled assumptions; for this reason, we do not attempt to compute a population threshold here.

In summary, previous attempts to calculate the neuronal pool size that could account for behavioral sensitivity (Massot et al. 2011) suffer from multiple incorrect assumptions. In fact, it has become increasingly clear that, even with measurements of pairwise correlations among neural responses and with *simultaneously recording* neural activity and behavior, it is very difficult to constrain quantitative comparisons of neural population sensitivity and behavior. This is because conclusions depend critically on both the detailed multi-dimensional covariance structure of population activity, as well as the type of decoder (Haefner et al. 2013; Moreno-Bote et al. 2014; Pitkow et al. 2015). Both of these properties are presently impossible to identify unambiguously using existing technologies.

### Correlations between neural activity and behavior

The functional significance of trial-by-trial correlations between neural activity and perceptual decisions, as typically quantified using choice probabilities (CPs), has been under debate. An early interpretation of choice-related activity is that it reflects the ‘bottom-up’ contribution of neural noise to perceptual decision-making processes (Britten et al. 1996; Shadlen and Newsome 2001), such that CPs could arise through feed-forward processing. More recently, there has been growing evidence that at least a substantial component of CP reflects ‘top-down’ signals (Cohen and Newsome 2008; Krug 2004; Nienborg and Cumming 2007; 2009; 2010; Zaidel et al. 2017; Yang et al 2016; Haefner et al. 2016; Wimmer et al 2015). Thus, fundamental issues regarding the origin of CPs remain unresolved.

Traditionally, CPs have been measured in cortex. An advantage of the vestibular system for studying choice-related activity is that the same basic forms of directional selectivity are seen at many levels of processing, from afferents to cortex (Laurens et al. 2017), unlike in other sensory systems. Thus, CPs can readily be measured in subcortical neurons using vestibular stimuli. Indeed, Liu et al (2013a) showed, for the first time, that VN neurons exhibit large CPs during a heading discrimination task. Furthermore, the inverse correlation between CPs and neuronal thresholds was steeper for VN neurons than cortical neurons (Liu et al 2013a).

The factors that drive expression of CPs in some cells, but not others, remain unclear. One hypothesis is that CPs appear when sensory signals are represented in an appropriate format for mediating the particular behavior. Our previous study (Liu et al. 2013a) provided support for this hypothesis by showing that CPs correlate with the degree to which VN/CN neurons represent translation (heading) without being confounded by orientation changes relative to gravity (as is the case for otolith afferents). In fact, when we found no significant CPs in otolith afferents (Yu et al. 2015), which all suffer from the gravito-inertial ambiguity (Angelaki et al. 2004), we reasoned that responses to the total gravito-inertial acceleration may be less useful for heading perception. In that case, one may argue that otolith afferent recordings cannot test the bottom-up origin of CPs because the otolith afferents do not really represent the task variable. In contrast, canal afferent responses are in the correct basic form to provide signals directly usable for rotation perception. Here, we recorded from canal afferents during rotation discrimination, and we find that their firing rates do not correlate with perceptual decisions regarding direction of rotation. Thus, these findings do not support a bottom-up origin of CPs in which variability in the afferents drives variability in percepts.

Further, we did not find significant choice-related activity in the VN during rotation discrimination. This result is surprising, given that translation-selective VN neurons show significant correlations with choice during heading discrimination (Liu et al. 2013a). The reasons for this difference between rotation and translation are unclear, and a number of factors might contribute. One possibility is that we may not have recorded from a subset of VN neurons that contribute to rotation perception, and that those neurons would show significant CPs. In fact, in this study, we identified proportionally much fewer neurons that were selectively responsive to rotation than in previous studies (Dickman and Angelaki 2002). One notable difference was that we used linear electrode arrays and off-line spike sorting, such that our sampling of neurons was less biased than in previous studies, which used single electrodes. Indeed, in the recordings described here, we found a surprisingly large proportion of bidirectional neurons. Such cells, which have been previously reported in the VN of reduced animal preparations (Type III classification of Duensing and Schaeffer 1958), have not been previously described in the VN of alert animals. Interestingly, the majority of these cells were characterized by low spontaneous firing rates, which may have resulted in them being excluded from traditional single electrode studies. Thus we cannot exclude the possibility that we would have observed significant CPs if we had sampled from a larger number of rotation-only cells, but this possibility does not seem very likely.

Additionally, the presence of significant CPs may depend on the structure of the task. In the present study, we used a coarse (left vs. right) discrimination task, whereas Liu et al. (2013a,b) used a fine heading discrimination task around a forward direction. It might be possible that neural variability in early stages of vestibular processing contributes more to variability of perceptual decisions for fine than coarse tasks. A third possibility might involve differences in the nature of the computations underlying estimation of rotation and translation. One such difference is that otolith afferent signals have a fundamental ambiguity between tilt and translation whereas canal afferent signals do not have a corresponding ambiguity. However, this explanation seems unlikely as it might have been thought to predict greater CPs for rotation than translation, which is opposite to what we found. Another difference lies in the commissural system; central canal and otolith pathways are fundamentally different (Uchino et al. 1997; 1999; 2001; Bai et al. 2002; Ogawa et al. 2000; Precht and Shimazu 1965; Precht et al. 1966; Shimazu and Precht 1966; Kasahara and Uchino 1974). At present, the reasons for the observed differences in CPs of VN neurons between translation and rotation perception remain fundamentally unclear.

Relevant to the origin of CPs is the extent of ‘shared variability’ among neurons, which is typically measured as correlated noise between pairs of neurons. Computational studies have shown that CPs depend on the structure of correlated noise in a population: without correlated noise, sizeable CPs are possible only with implausibly small neural pool sizes (Shadlen et al. 1996; Haefner et al. 2013; Pitkow et al 2015; Gu et al 2014). For moderate to large populations, the presence of significant CPs requires nonzero correlated noise, but the converse is not true. The absence of significant CPs during rotation discrimination might have been explainable if there were no noise correlations during rotation, but we found robust noise correlations during rotation discrimination with a relationship between noise and signal correlations that is roughly similar to that observed previously for heading (translation) discrimination (Liu et al. 2013a). Thus, the correlated noise that we measured during rotation discrimination in VN appears to reflect signals that are not related to choice.

### Conclusion

The present findings show that neurons in the peripheral vestibular system have sensitivity approaching that of the animal’s behavior – and this has been a frequent finding in other sensory domains (e.g., Britten et al 1992; Chen et al 2013a; Cohen & Newsome 2009; Gu et al 2007; Uka and DeAngelis 2003, 2006). *So, why isn’t perception, which presumably arises from large populations of neurons, much more sensitive than single cells?* The fact that single canal afferents have thresholds that are similar to behavioral thresholds is remarkable, given that there are on the order of ~3,000 afferents innervating the same vestibular organ (Wilson and Melvill Jones 1979). Pairwise noise correlations are small in canal afferents (Yu et al. 2014); thus, either correlations of higher order strongly limit information in the population or decoding is extremely suboptimal in the context of our task. Finding the answers to these questions will be fundamental for understanding sensory perception, and the vestibular system provides an excellent model system for continuing investigations.

## Notes

Work supported by NIH DC004260, DC004260-18 Diversity Supplement, and NIH IMSD 5R25 GM056929 Initiative for Maximizing Student Diversity.

